# *Hey2* restricts cardiac progenitor addition to the developing heart

**DOI:** 10.1101/199638

**Authors:** Natalie Gibb, Savo Lazic, Ashish R. Deshwar, Xuefei Yuan, Michael D. Wilson, Ian C. Scott

**Affiliations:** Program in Developmental and Stem Cell Biology, The Hospital for Sick Children, Toronto, Ontario, M5G 0A4, Canada; Program in Genetics and Genome Biology, The Hospital for Sick Children, Toronto, Ontario, M5G 0A4, Canada; Department of Molecular Genetics, University of Toronto, Canada; Ted Rogers Centre for Heart Research, Toronto, Canada; Heart and Stroke Richard Lewar Centres of Excellence in Cardiovascular Research, Toronto, Canada

## Abstract

A key event in vertebrate heart development is the timely addition of second heart field (SHF) progenitor cells to the poles of the heart tube. This accretion process must occur to the proper extent to prevent a spectrum of congenital heart defects (CHDs). However, the factors that regulate this critical process are poorly understood. Here we demonstrate that Hey2, a bHLH transcriptional repressor, restricts SHF progenitor accretion to the zebrafish heart. *hey2* expression demarcated a distinct domain within the cardiac progenitor population. In the absence of Hey2 function an increase in myocardial cell number and SHF progenitors was observed. We found that Hey2 limited proliferation of SHF-derived cardiomyocytes in a cell-autonomous manner, prior to heart tube formation, and further restricted the developmental window over which SHF progenitors were deployed to the heart. Taken together, our data suggests a role for Hey2 in controlling the proliferative capacity and cardiac contribution of late-differentiating cardiac progenitors.

## INTRODUCTION

Cardiac development is regulated by the activity of concerted signaling, transcriptional and morphogenetic events. Subtle perturbations in these processes, either genetic or environmental, can lead to congenital heart defects (CHD), the most common class of congenital anomalies. It is now evident that the vertebrate heart is built from two populations of progenitor cells, termed the first heart field (FHF) and second heart field (SHF), which contribute to the heart in two successive windows of differentiation. Cells of the FHF differentiate in an initial wave of cardiogenesis, resulting in formation of the linear heart tube. Over a well-defined developmental window, multi-potent, late-differentiating progenitors of the SHF migrate into the poles of the heart tube to extensively remodel and add structure to the heart (Cai et al., 2003; Hutson et al., 2010; Kelly, 2012; van den Berg et al., 2009). It remains under debate however whether these populations of cardiac progenitor cells (CPCs) represent distinct populations with unique molecular signatures, or whether they exist as one population with a gradient in timing for deployment to the heart (Abu-Issa et al., 2004; Ivanovitch et al., 2017; Moorman et al., 2007).

In zebrafish, several fate-mapping studies have found that SHF progenitors give rise to the distal portion of the ventricular myocardium and smooth muscle of the outflow tract (OFT; (Guner-Ataman et al., 2013; Hami et al., 2011; Zeng and Yelon, 2014; Zhou et al., 2011). Embryological manipulations in chick and SHF-restricted mutation of CHD-associated genes in the mouse have firmly established that defects in SHF development are a major contributor to CHD (Cai et al., 2003; Prall et al., 2007; Ward et al., 2005). As a key driver of cardiac morphogenesis, the balance of proliferation, “stemness” and cardiac differentiation events in the SHF progenitor pool must be tightly regulated. Several signaling pathways have been implicated in the development of the SHF (Li et al., 2016; Mandal et al., 2017; Prall et al., 2007; Ryckebusch et al., 2008; Sirbu et al., 2008; Tirosh-Finkel et al., 2010; Zhao et al., 2014). Notable amongst these are fibroblast growth factor (FGF) and retinoic acid (RA) signaling. The permissive signals from RA towards FGF signaling create a mutual opposition for regulating the specification and differentiation of cardiac progenitor populations (Ilagan et al., 2006; Park et al., 2008; Rochais et al., 2009; Sorrell and Waxman, 2011; Witzel et al., 2012). Transcription factors including Islet1 (Isl1), Nkx2.5 and Fosl2 are essential for maintaining the SHF progenitor pool (Cai et al., 2003; de Pater et al., 2009), yet the full make-up of the transcriptional network required to precisely regulate the behavior of the late-differentiating cardiac progenitor population remains unclear.

Hey2 is a member of the Hairy/Enhancer of split basic Helix-Loop-Helix (bHLH) subfamily, which act as transcriptional repressors during embryonic development (Davis and Turner, 2001). Zebrafish *hey* genes exhibit more restricted expression patterns compared to mammals, with *hey2* being the only family member detectably expressed in the heart (Winkler et al., 2003). Studies in mice have shown that a disruption in *hey2* can lead to ventricular septal defects (VSD) as well as other CHDs and cardiomyopathy (Donovan et al., 2002; Sakata et al., 2002). Mutation in zebrafish *hey2* leads to a localized defect of the aorta resembling human aortic coarctation (Weinstein et al., 1995; Zhong et al., 2000), with Hey2 having a key role in specifying arterial versus venous cell fates (Hermkens et al., 2015; Zhong et al., 2001; Zhong et al., 2000). Of note, Hey2 has been suggested to regulate growth of the heart via restraining cardiomyocyte proliferation (Jia et al., 2007).

Based on computational approaches, Hey2 has recently been predicted to be a key regulator of human cardiac development (Gerrard et al., 2016). Given the evidence linking Hey2 function with CHD-associated defects in various model systems, and the association between CHD and SHF progenitors, we hypothesized that *hey2* may have a role in regulating the late-differentiating SHF progenitor pool. Using the zebrafish we were able to discover a distinct domain of *hey2* expression localized to regions adjacent to the myocardium, suggestive of a function for Hey2 in the SHF. Analysis of a novel null *hey2* mutant allele revealed increased myocardial cell number, with an apparent increase in the size of the SHF progenitor pool at multiple stages of cardiac development. Temporal analysis demonstrated that progenitor cells underwent increased proliferation prior to, but not following, addition to the heart in the absence of Hey2 function. This led to both more robust and extended late addition of cardiac progenitors to the heart, with *hey2* acting in a cell-autonomous manner in this context. Taken together, these results suggest that *hey2* acts as a key brake on the proliferative capacity and deployment of SHF progenitors to the vertebrate heart.

## EXPERIMENTAL PROCEDURES

### Zebrafish husbandry and transgenic lines

Adult zebrafish were maintained as per Canadian Council on Animal Care (CCAC) and The Hospital for Sick Children Animal Services (LAS) guidelines. Zebrafish embryos were grown at 28.5°C in embryo medium as previously described (Westerfield, 1993). The following transgenic lines were used: *Tg*(*myl7:EGFP)*^*twu34*^, (Huang et al., 2003), *Tg(nkx2.5:ZsYellow)*^*fb7*^ (Zhou et al., 2011), Tg(*myl7:nlsKikGR)*^*hsc6*^ (Lazic and Scott, 2011), *Tg(myl7:nlsDsRedExpress)*^*hsc4*^ (Lou et al., 2011), *Tg(gata1:DsRed)*^*sd2*^ (Traver et al., 2003) and *Tg(myl7:mCherry-RAS)*^*sd21*^ (Yoruk et al., 2012). The *hey2*^*hsc25*^ mutant line was generated using CRISPR/Cas9 genome editing technology as previously described (Jao et al, 2013). Primers TAGGCCAGAAAGAAGCGGAGAG and AAACCTCTCCGCTTCTTTCTGG were annealed together and ligated into the pT7-gRNA vector digested with BsmBI to create sgRNA for *hey2*. An 8bp deletion allele (starting at nucleotide 151 within exon2) resulting in a premature stop codon at amino acid 53 was isolated (Fig. 3A). An additional allele harboring a 5bp deletion (starting at nucleotide 152 with a premature stop codon at amino acid 55) was isolated showing an equivalent phenotype (data not shown). The *hey2* enhancer transgenic *epiCon21:EGFP*^*hsc28*^, containing an epigenetically-conserved open chromatin region (epiCon21), was identified from comparative epigenetic analysis (XY, MDW and ICS, manuscript in preparation). The 626bp enhancer sequence, located 24kb upstream of the *hey2* locus (supplemental Fig. 1A), was amplified from zebrafish genomic DNA (FP 5’ – CAAATCCCCTGACCTCTGCTTTGAG – 3’ RP 5’ – GACACACAGTGACATGTCCTATTGCG – 3’) and cloned into the enhancer detection vector *E1b-Tol2-GFP-gateway* (Addgene 37846 (Li et al., 2010). To generate *Tg(epiCon21:EGFP)*, 25ng of *E1b-Tol2-GFP-gateway* plasmid carrying the epiCon21 enhancer was injected into wild type embryos at the one-cell stage with 150ng *Tol2* mRNA. Four independent germline carriers were identified, which demonstrated indistinguishable patterns of GFP expression. The *epiCon21:EGFP*^*hsc28*^ was maintained and used for all experiments. Generation of *Tg(hey2V5*^*hsc27*^) was performed as previously described (Burg et al., 2016). nCas9n mRNA was injected at a concentration of 150pg together with 30pg of *hey2*gRNA into the yolks of 1-cell embryos. V5 tagging oligo (5’ – AGTCATGGCCAGAAAGAAGCGGCAAGCCTATCCCAAACCCTCTGCTGGGCCTGGACTCCACAGGAGAGGGGTAATTCATATT – 3’) was diluted to 25μg/μl and 1nl was injected into the yolks immediately after the RNA injections.

**Figure 1.**
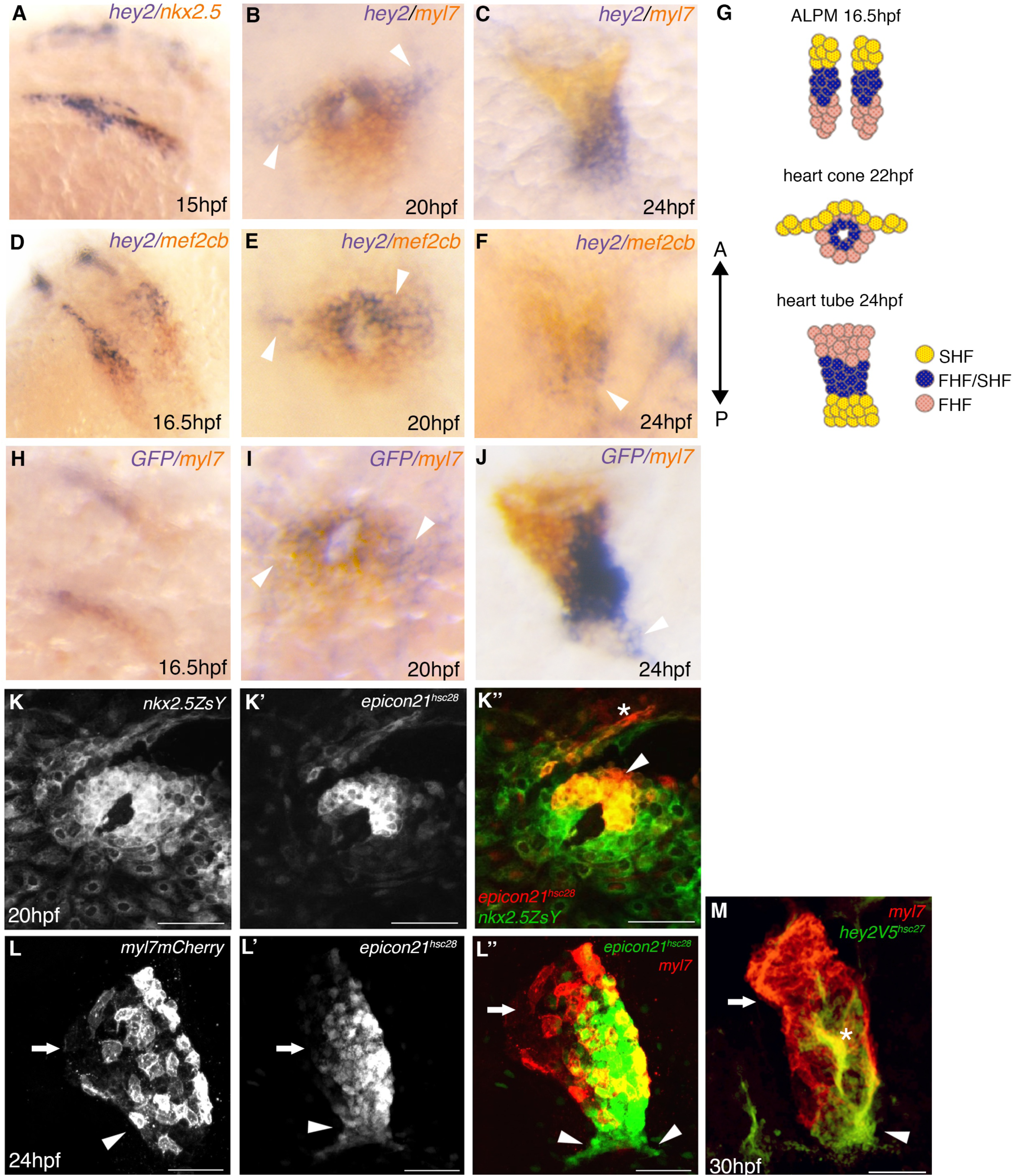
*hey2* resides in the late-differentiating progenitor population. (A-F) Double RNA *in situ* hybridization analysis of *hey2* (blue) against *nkx2.5*, *myl7* and *mef2cb* (orange) in wild type embryos from 16.5 hpf to 24 hpf. (G) Schematic representation of FHF and SHF progenitor localization and movement patterns from 16.5 to 24 hpf. (H-J) Double RNA *in situ* hybridization analysis of *Tg(epiCon21:EGFP)*^*hsc28*^ (blue) an*d myl7* (orange) at 15 hpf (H), 20 hpf (I) and 24 hpf (J). (K-L) Immunofluorescence showing *Tg(epiCon21:EGFP)*^*hsc28*^ enhancer expression against *Tg(nkx2.5:ZsYellow)* at 20 hpf (K-K’’) and 24 hpf (L-L’’). (M) Immunofluorescence of *Tg(hey2-V5)*^*hsc27*^ internal epitope tagging with *Tg(myl7:mCherry-RAS)* at 30 hpf comparing Hey2 expression (green) with Myl7 (red). Scale bars 50μm. Asterisk labels pharyngeal mesoderm. Arrows denote the atrium and arrowheads mark the ventricle.

**Figure 3.**
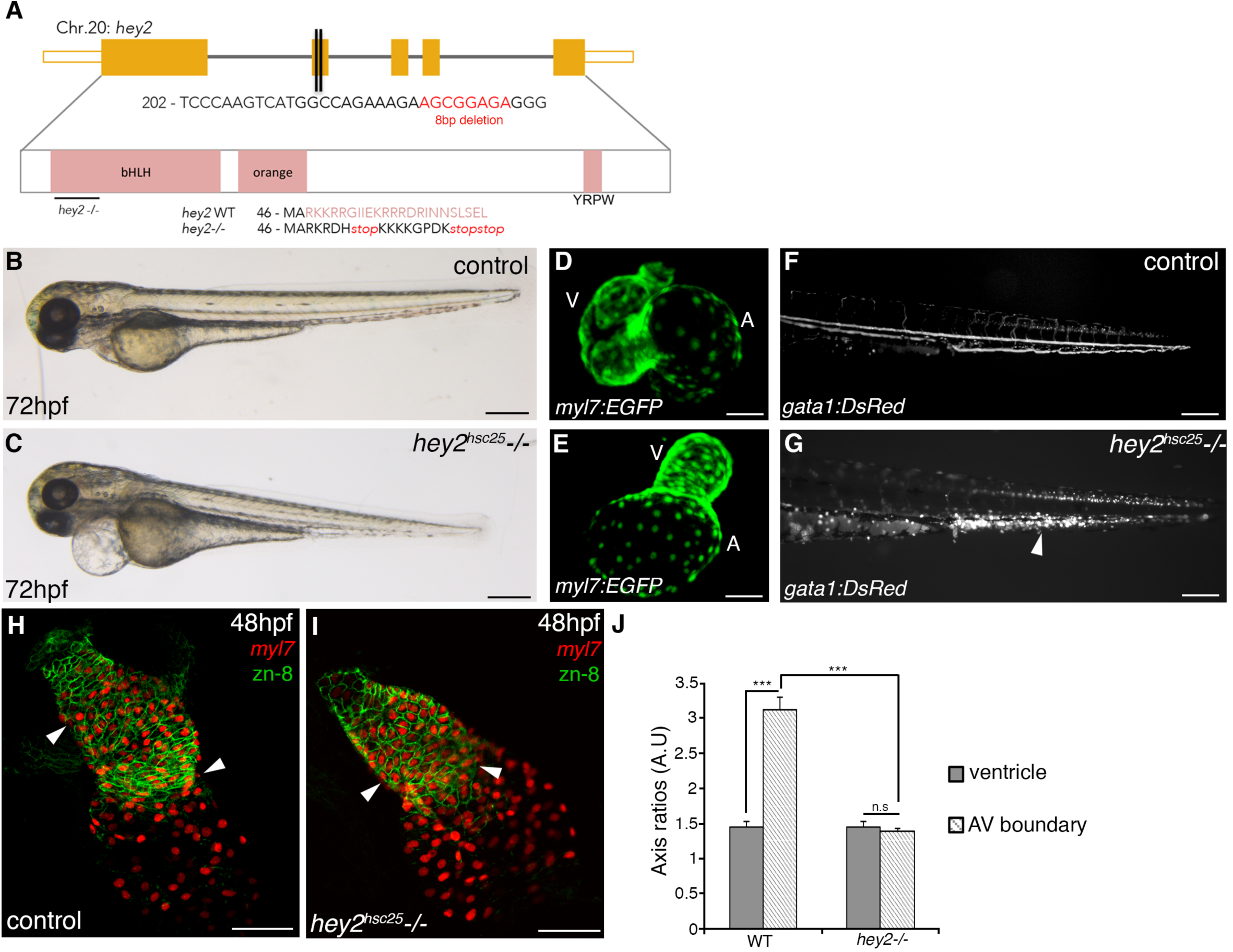
Cardiovascular defects are observed in the absence of Hey2 function. (A) Schematic representation of *hey2* null-mutant generated through CRISPR/Cas9 mediated genome-editing. Red lettering shows 8bp deleted sequence. Protein sequence shows production of premature stop codon at the beginning of exon 2. (B, C) Bright-field images of a sibling control and a *hey2*^*hsc25*^ mutant embryo at 72hpf. (D, E) Confocal images of *Tg(myl7:EGFP)* hearts in control (D) and *hey2*^hsc25^ *-/-* (E) embryos at 72 hpf. (F, G) Fluorescent images of *Tg(gata1:DsRed)* showing normal blood flow in controls at 72 hpf (F) and lack of blood flow leading to coagulated blood cells in *hey2*^hsc25^ *-/-* (G, arrowhead). (H-I) Confocal imaging of *Tg(myl7:nlsDsRedExpress)* embryos co-stained with zn-8 to display cell number (DsRed) and cell membrane (zn-8) formation in control (H) and *hey2*^*hsc25*^ mutant embryos (I) at 48 hpf. (J) Bar graph showing axis ratios of cardiomyocytes taken from the ventricle and the AV boundary between control and *hey2*^*hsc25*^ mutant embryos. N=3, n=10 per condition. Error bar mean±s.e.m; *** p<0.001; n.s, not significant. Scale bar 50μm (B-G) and 100μm (H and I).

### Morpholinos

Morpholino oligos were purchased from Genetools (Oregon, USA). A morpholino targeting the translation start site of *hey2* (ATG *hey2*: 5’ – TGCTGTCCTCACAGGGCCGCTT<u>CAT</u> - 3’) was used throughout this study. The *hey2* morpholino (5’ – CGCGCAGGTACAGACACCAAAAACT - 3’) previously described (Jia et al., 2007) was used to test specificity, as both morpholinos share no sequence overlap. Injection of 1ng of ATG *hey2* morpholino at the one-cell stage yielded a consistent heart phenotype.

### Standard RNA *in situ* hybridization

Standard RNA *in situ* hybridization was performed as previously described (Thisse and Thisse, 2008). The complete coding sequence of *hey2* (ZDB-GENE-000526-1) was PCR amplified (sense 5’–ATGAAGCGGCCCTGTGAGGACAGC, antisense 5’– TTAAAACGCTCCCACTTCAGTTCC) and used as a riboprobe template. GFP riboprobe sequence was cloned into pGEM-Teasy and transcribed per standard techniques. Previously described riboprobes were additionally used: *hand2* (ZDB-GENE-000511-1), *myl7* (ZDB-GENE-991019-3), *nkx2.5* (ZDB-GENE-980526-321)*, mef2cb* (ZDB-GENE-040901-7), *amhc* (ZDB-GENE-031112-1), *vmhc* (ZDB-GENE-991123-5), *tbx1* (ZDB-GENE-030805-5), *ltbp3* (ZDBGENE-060526-130), *bmp4* (ZDB-GENE-980528-2059) and *tbx2b* (ZDB-GENE-990726-27) (Chen and Fishman, 1996; Lazic and Scott, 2011; Yelon et al., 1999). DIG and Fluorescein-labeled probes were made using a RNA Labeling Kit (Roche).

### Quantitative RT-PCR

Quantitative real-time PCR was performed using the Roche LightCycler 480 with Platinum SYBR green master mix used as per manufactures instructions (ThermoFisher Scientific 11733038). Primers used are as follows: *hey2* forward 5’ GTGGCTCACCTACAACGACA 3’, reverse 5’ CCAACTTGGCAGATCCCTGT 3’ *mef2cb* forward 5’ CAGCCCAGAGTCAAAGGACA 3’, reverse 5’ AGGGCACAGCACATATCCTC 3’ and *nkx2.5* forward 5’ TCTCTCTTCAGCGAAGACCT 3’ reverse 5’ CTAGGAAGTTCTTCGCGTAA 3’. Previously described primers were used for quantification of *β-actin* (Tang et al., 2007); *ltbp3* (Zhou et al. 2011); *tbx1* (Zhang et al., 2006) and *amhc* (Jia, et al 2007) transcript levels.

### Small molecule treatments

The FGF receptor inhibitor SU5402 (Tocris 3300) was used at a concentration of 10μM from 16.5 to 20 hours post-fertilization (hpf) or from 19 to 24 hpf. BMP and Notch signaling inhibitors dorsomorphin (Tocris 3093) and DAPT (Tocris 2634/10), respectively, were used at a concentration of 10μM and 50μM between 16.5 and 20 hpf. Retinoic Acid (Sigma R2625) was added at a concentration of 0.1μM to dechorionated embryos at 5.3 hpf (50% epiboly) for 1 hour. All compounds were diluted into 1% DMSO in embryo medium. Vehicle controls were treated with 1% DMSO. Incubations were performed at 28°C.

### Imaging

Bright-field images were taken using a Zeiss AXIO Zoom V16. RNA *in situ* hybridization images were captured using a Leica M205FA microscope with the LAS V6 software package. Immunofluorescence (IF) confocal images were taken with a Nikon A1R laser scanning confocal microscope.

### Immunofluorescence, DAF-2DA staining and cell counting

Whole-mount IF was carried out as previously described (Alexander et al., 1998). Primary antibodies used were: α-MYH6 supernatant 1:10 (DSHB, S46); α-MHC supernatant 1:10 (DSHB, MF20); α-MEF-2 (C21) 1:250 (Santa Cruz sc-313); α- RCFP 1:400 (Clontech 632475); α-Neurolin (cd-166) supernatant 1:10 (DSHB, ZN- 8); α-DsRed 1:200 (Clontech 632496); α-V5 1:500 (ThermoFisher Scientific R960- 25) and α-GFP 1:1000 (Torrey Pines Biolabs). Smooth muscle of the bulbus arteriosus was visualized using the NO indicator DAF- 2DA (Sigma D2813) as previously described (Grimes et al., 2006). Cardiomyocyte nuclei of *myl7:nlsDsRedExpress* transgenic embryos were counted following immunostaining. Embryonic hearts were dissected and flat-mounted prior to confocal imaging.

### Photoconversion and cell addition analysis

Photoconversion on *myl7:nlsKikGR* embryos was carried out as previously described (Lazic and Scott, 2011) using the UV channel on a Zeiss Axio zoom V16 microscope. Images were captured using a Nikon A1R laser scanning confocal. For mounting, embryos were fixed in 4% PFA for 20 minutes and washed three times in PBS. Embryos were agitated in 5% saponin/PBS 0.5%Tx-100 followed by dehydration to 75% glycerol/PBS and left overnight at 4°C. Hearts were dissected and flat mounted prior to imaging.

### EdU incorporation

EdU incorporation assays were performed as previously described (Zeng and Yelon, 2014). Embryos were incubated in 10mM EdU at 16, 22 and 24 hpf on ice for 30 minutes. A Click-iT imaging kit (Invitrogen) was used to visualize EdU incorporation. *Tg(nkx2.5:ZsYellow)* expression was detected using α-RCFP (Clontech 632475) with Alexa Fluor 488-conjugated goat anti-rabbit secondary. A proliferation index was calculated based on cells positive for both EdU (red) and ZsYellow (green) staining.

### Statistical analysis

Excel software was used to perform student t-tests with two-tail distribution. Graphs display mean±s.e.m unless otherwise stated. Box plot graphs were prepared using BoxPlotR web-tool (http://boxplot.tyerslab.com).

## RESULTS

### *hey2* expression marks a subset of cardiac progenitor cells

To further examine the role of Hey2 in cardiac development, we first carried out a detailed analysis of *hey2* expression during key stages of cardiogenesis, spanning early cardiac specification to the formation of the linear heart tube, using whole-mount RNA *in situ* hybridization. Interestingly, *hey2* transcripts were found to localize anteromedial to those of *nkx2.5* and *mef2cb* in the anterior lateral plate mesoderm (ALPM) at 16.5 hours post-fertilization (hpf, Fig. 1A and D, respectively). At 20 hpf, when the primitive heart is organized into a cone of differentiating cardiac cells (Yelon et al., 1999), *hey2* expression was again evident anteromedial to that of *myl7* and *mef2cb* (Fig. 1B and E). We further observed a domain of *hey2* expression lateral to the heart cone, in the region of the pharyngeal mesoderm (Fig. 1B and E, white arrowheads). Following formation of the linear heart tube at 24 hpf, *hey2* transcripts were detectable both within and extending from the distal portion of the ventricle, a region occupied by *mef2cb*-positive cells of the presumptive SHF (Fig. 1C and F; (Lazic and Scott, 2011). These results, as summarized in Figure 1G, suggest that *hey2* is an early marker of the late-differentiating progenitor population, as it is expressed in a manner consistent with regions shown to contain SHF progenitors (Guner-Ataman et al., 2013; Hami et al., 2011).

To further dissect the expression of *hey2* with respect to the late-differentiating progenitor population, we pursued both isolation of key *hey2* regulatory regions and tagging of the endogenous *hey2* coding sequence. Epigenetic analysis of early zebrafish cardiogenesis identified an enhancer (epiCon21) located 24kb upstream of *hey2* that shares an open chromatin signature between zebrafish, mouse and humans (supplemental Fig. 1A; XY, MDW and ICS, manuscript in preparation). Stable *Tg(epiCon21:EGFP)* transgenic animals were made in which this enhancer drove GFP expression. Following RNA *in situ*, analysis of *Tg(epiCon21:EGFP)* and *myl7* expression at 16.5, 20 and 24 hpf showed that the transgenic faithfully recapitulated the endogenous *hey2* gene expression pattern (Fig. 1 compare H-J with A-C). Higher resolution analysis revealed *Tg(epiCon21:EGFP)*^*hsc28*^ expression at 20 hpf restricted to the anteromedial region of the *Tg(nkx2.5ZsYellow)*-positive heart cone, as was observed by RNA *in situ* hybridization (Fig. 1 B and K-K’’). The higher resolution afforded by fluorescent immunohistochemistry further indicated *Tg(epiCon21:EGFP)* co-expression with *Tg(nkx2.5ZsYellow)* in the pharyngeal mesoderm (Fig. 1K’’, asterisk; (Paffett-Lugassy et al., 2013). A further domain of *Tg(epiCon21:EGFP)* expression was also observed immediately anterior to the heart cone, with these cells having no detectable *Tg(nkx2.5ZsYellow)* expression (Fig. K’’ arrowhead), matching the position of recently described *isl2b*-positive SHF cells (Witzel et al., 2017). By 24 hpf, *Tg(epiCon21:EGFP)*^*hsc28*^ shows clear expression within the posterior ventricular region, with the most posterior domain being negative for *myl7* expression (Fig. 1L- L’’ arrowheads indicate ventricular portion and arrows indicate atrial portion of the heart tube). Analysis of the expression of *Tg(epiCon21:EGFP)*^*hsc28*^ against that of *Tg(myl7:mCherry)* until 94 hpf demonstrated specificity to the ventricular myocardium and OFT, with an absence of detectable expression in the atrium and atrio-ventricular canal (supplemental Fig. 1B-F).

To conclusively follow endogenous *hey2* expression, we further used CRISPR/Cas9 genome editing to place an internal V5 epitope tag into the *hey2* locus (supplemental Fig. 1G; Burg et al, 2016). *T*g(*hey2-V5)*^*hsc27*^ embryos were viable, demonstrating that this allele was functional. Antibody staining versus V5 at 30 hpf showed V5-Hey2 localization to the posterior ventricular region of the heart tube (Fig. 1M; arrowhead ventricle; arrow atrium), with a portion of CMs expressing both *myl7:mCherry* and V*5* (Fig. 1M; asterisk). This expression data replicates both endogenous expression of *hey2* (Fig. 1C) as well as enhancer expression of *hey2* (Fig. 1J and L’’). Taken together, expression analysis revealed subsets of *hey2*-positive cells between 15 and 20hpf which 1) only express *hey2*; 2) co-express both *hey2* and *nkx2.5* in the cardiac cone; and 3) co-express *hey2* and *nkx2.5* in presumptive cardiac progenitors within the pharyngeal mesoderm.

### Opposing effects of FGF and RA signaling on *hey2* expression suggest a link to SHF progenitors

To further explore the link between Hey2 and the late-differentiating progenitor population, we reasoned that *hey2* expression should be affected by modulation of signaling pathways that have been implicated in regulating SHF progenitor development. Both FGF and retinoic acid (RA) signaling have been shown to play opposing roles during cardiac specification, diversifying the heart fields within the ALPM (Ryckebusch et al., 2008; Sirbu et al., 2008; Waxman et al., 2008). Subsequent to this, FGF signaling is required within the late-differentiating SHF progenitors for proper OFT development and maintenance of progenitor proliferation and survival (Ilagan et al., 2006; Park et al., 2008; Zeng and Yelon, 2014). Previous reports identified *hey2* as a target of both FGF and RA signaling, with RA acting via FGF (Feng et al., 2010; Sorrell and Waxman, 2011).

By modulating FGF signaling between 16.5 hpf and 20 hpf using a well characterized inhibitor, SU5402, we noted the expected reduction in ventricular cardiomyocyte differentiation but normal atrial differentiation, as shown by *vmhc* and *amhc* expression (supplemental Fig. 2A and B). SU5402 treatment between 16.5 hpf and 20 hpf further resulted in reduced cardiac *hey2* expression within the heart cone (Fig. 2A and B), with a coincident loss of detectable expression of the SHF progenitor marker *mef2cb* (supplemental Fig. 2C and D, (Lazic and Scott, 2011). Later addition of SU5402, between 19 and 24 hpf, resulted in a loss of cardiac *hey2* expression, yet no overt change to cardiac *myl7* or neural *hey2* expression (Fig. 2C and D; arrowhead cardiac, asterisk neural). While this result showed coincident loss of SHF-associated progenitors and *hey2* expression, we could not preclude that the absence of *hey2* simply reflected its association with a ventricular fate.

**Figure 2.**
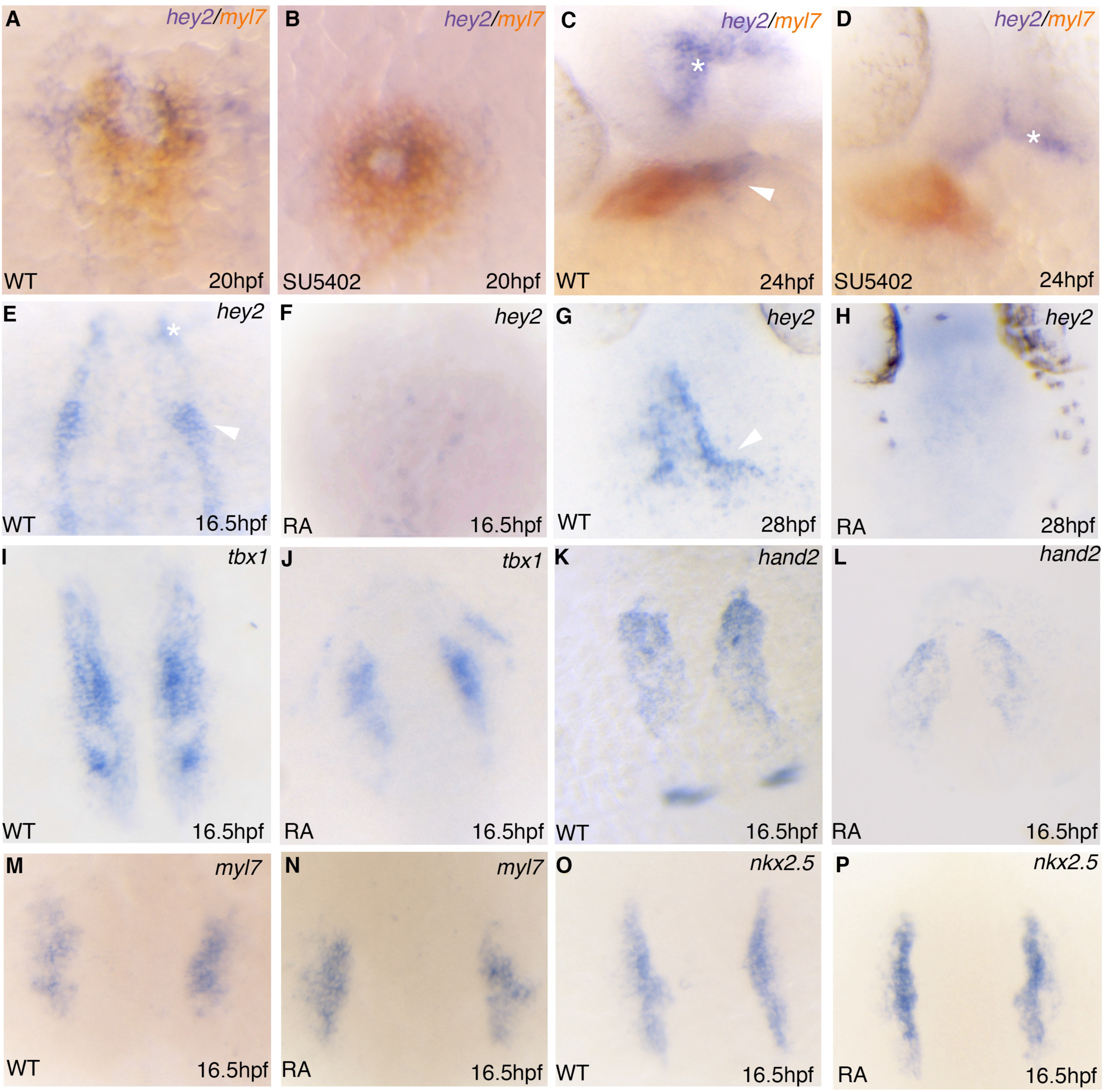
Opposing effects of FGF and RA signaling regulate *hey2* expression. (A-D) Double RNA *in situ* hybridization analysis of *hey2* (blue) and *myl7* (orange) expression on embryos treated with 10μM SU5402 from 16.5 – 20 hpf (A and B) and from 19 – 24 hpf (C and D). (E-P) RNA *in situ* hybridization analysis for *hey2* (E-H), *tbx1* (I and J), *hand2* (K and L), *myl7* (M and N) and *nkx2.5* (O and P) following treatment with RA at 0.1μM for 1 hour at 50% epiboly. Note down regulation in *tbx1* and *hand2* but no change in *myl7* and *nkx2.5*. Asterisks depict neural expression, arrowheads cardiac expression.

To further examine the potential relationship between *hey2* expression and SHF progenitors, we next modulated RA activity. The effect of RA on SHF development was studied via use of a photoconversion assay in *myl7:nlsKikGR* embryos via quantifying later (SHF-derived) addition of cardiomyocytes to the heart tube after 24 hpf (Lazic and Scott, 2011). Addition of RA at 4 hpf (50% epiboly) inhibited myocardial accretion between 24 and 48 hpf as shown by a reduction in green-only cardiomyocytes as well as a significant loss of ventricular CM number (supplemental Fig. 2E-H). This highlights the inhibitory effect of RA on the late-differentiating progenitor population. In keeping with this result, *hey2* expression was undetectable under the same conditions of exogenous RA treatment at both 16.5 and 28 hpf (Fig. 2E-H). Loss of *hey2* expression following RA treatment was accompanied by a reduction in expression o*f tbx1* and *hand2* (Fig. 2I-L) as well as a loss of *mef2cb* expression (supplementary Fig. 2I and J), all of which are associated with SHF development. In contrast, no appreciable effects on *nkx2.5* or *myl7* expression was observed at either 16.5 hpf (Fig. 2M-P) or 28 hpf (supplemental Fig. 2K and L), suggesting comparatively normal FHF progenitor development. In contrast to FGF and RA signaling, inhibition of BMP and Notch pathways had no detectable effect on *hey2* expression by 20 hpf (supplemental Fig. 2M and N). While RA treatment affected both ventricular and SHF cell fate at 28 hpf, the early effect on expression of *hey2* and other SHF markers suggested a link between *hey2* and the late-differentiating population.

### Cardiovascular defects in the absence of *hey2* function

As expression of *hey2* was suggestive of a role in early cardiac development, prior to heart tube assembly, we next investigated the effect loss of Hey2 function. Previous reports demonstrated the consequences of knocking down *hey2* gene function using either morpholinos (MOs) or with existing *hey2* mutants: an ENU mutagenesis allele (*grl*^*m145*^ (Weinstein et al., 1995; Zhong et al., 2000) and in mosaic TALEN-injected embryos (Hermkens et al., 2015). However, overexpression experiments had suggested that the *grl*^*m145*^ allele, which encodes a Hey2 protein with a 44aa C-terminal extension, retains some function (Jia et al., 2007; Zhong et al., 2000). With both this and recent controversy regarding use of antisense morpholinos to assess gene function in zebrafish (Kok et al., 2015) in mind, we generated a novel predicted null mutation in *hey2* using CRISPR/Cas9-mediated genome editing. By targeting the *hey2* transcript within exon2, we generated a mutant with an 8bp deletion producing a premature stop codon at amino acid 53, effectively deleting the bHLH domain (Fig. 3A). Both *hey2*^*hsc25*^ mutant and morphant phenotypes become apparent by 48 hpf with mutant embryos displaying a non-looped heart with a reduction in heart rate when compared to controls (supplemental Fig. 3A-D). At 72 hpf, *hey2* mutant embryos exhibited pericardial edemas accompanied by an enlarged atrium, evident by a significant increase in *amhc* gene expression as well as a truncated ventricle (Fig. 3B-E, supplemental Fig. 3E-G). As in *grl*^*m145*^ mutants (Zhong et al., 2001), *hey2*^*hsc25*^ mutants demonstrated a blockage at the aortic bifurcation preventing blood flow to the trunk, evident by the slow movement of *gata1+* cells in the trunk vasculature (Fig. 3F and G, arrowhead). As *hey2* loss-of-function demonstrated a late morphological phenotypic onset, early developmental stages were analyzed using *hey2* morpholino to facilitate these studies.

### Loss of Hey2 affects cardiac function and maturation

Given the irregular morphology of *hey2* mutant hearts, we next investigated cardiomyocyte-intrinsic defects. The function of the heart is highly dependent on cardiomyocyte morphology, specifically cell shape (Auman et al., 2007). Cardiomyocytes initially have a uniform cuboidal characteristic that is altered as ventricular chamber formation proceeds by both blood flow and cardiac contractility (Manasek, 1981; Manasek et al., 1972; Taber, 2006). To examine potential defects in cardiac maturation, the *hey2*^*hsc25*^,allele was crossed into a *Tg(myl7:nlsDsRedExpress)* background. IF staining using antibodies against DsRed and zn-8 was used to visualize cardiomyocyte cell nuclei and cell membranes, respectively. As compared to sibling controls, in *hey2*^*hsc25*^mutants cardiomyocytes failed to initiate cellular elongation near the atrio-ventricular canal (AVC) (Fig. 3H and I, arrowheads). Instead, cell shape remained uniform throughout the ventricle. This was quantified through axis ratio measurements, which showed a significant difference in cell shape at the AV boundary in *hey2*^*hsc25*^ mutants compared to control embryos (Fig. 3J). To further characterize the defects observed within the AVC of *hey2*^*hsc25*^mutants, we performed RNA *in situ* hybridization. Upon AVC differentiation, *tbx2b* and *bmp4* expression become restricted to the AVC, with no transcript detected in the chamber myocardium (Rutenberg et al., 2006). We found that in *hey2*^*hsc25*^mutants *bmp4* and *tbx2b* transcripts remained distributed in the chambers, with no characteristic restriction to the AVC, a phenomenon also reported in mouse embryos misexpressing *hey2* (supplemental Fig. 3H-K; (Kokubo et al., 2007). These results suggest a function for Hey2 in proper AVC development and cardiac maturation.

### Expanded cardiomyocyte number in *hey2* mutants

Based on the expression of *hey2* prior to 24 hpf, we next analyzed the structure of the heart tube between wild type and *hey2* MO injected embryos. At 26 hpf, *hey2* morphants had an irregularly shaped heart that had failed to elongate (Fig. 4A-D), with an expansion in expression of terminal differentiation markers *myl7* and *tnnt2* at the poles of the heart (Fig. 4B and D, arrowheads). Given this observation, we counted CM number in wild type and *hey2* morphant *Tg(myl7nls:DsRedExpress)* embryos. Whereas control embryos contained 136.8±6 (mean±s.e.m, *n*=5) CMs at 24 hpf, *hey2* morphant embryos contained a significantly greater number (170±3.6, mean±s.e.m, *n*=5; Fig. 4E-G). As the heart develops, the number of CMs increases significantly between 24 and 48 hpf (de Pater et al, 2009). We therefore next determined the effect of *hey2* loss on 48 hpf CM number. As observed at 24 hpf, there was a significant increase in *myl7:nlsDsRedExpress*-positive CM nuclei in the absence of *hey2* as compared to controls (238.7±2.4 in *hey2*^*hsc25*^ mutants; 183±1.5 in controls, mean±s.e.m, *n*=6; Fig. 4F-H). These results highlight that in the absence of *hey2*, an elevated addition of CMs to the heart is detectable as early as 24 hpf.

**Figure 4.**
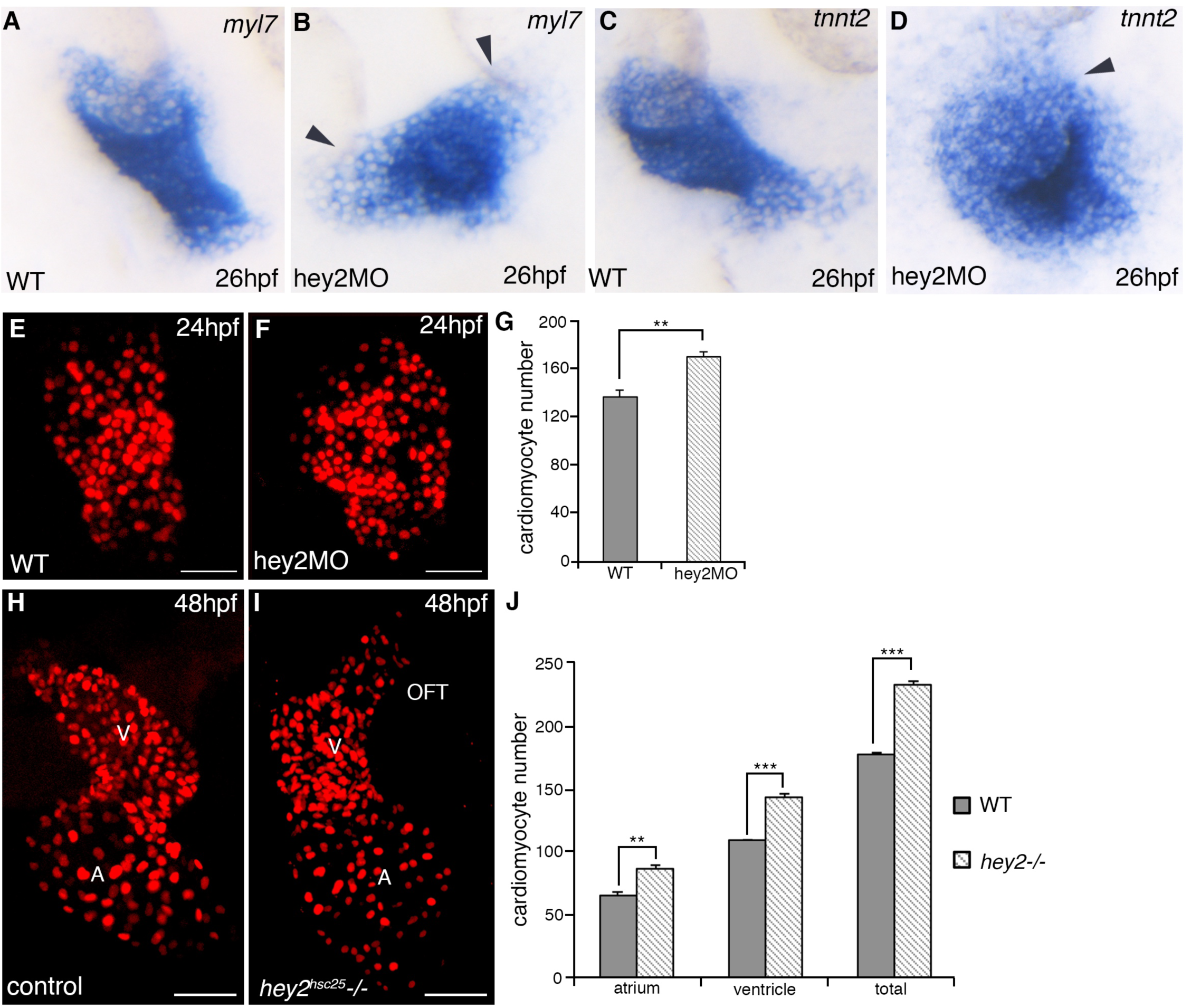
The absence of Hey2 results in inappropriate cardiomyocyte number. (A-D) RNA *in situ* hybridization analysis for *myl7* (A, B) and *tnnt2* (C, D) in control and hey2MO embryos. (E, F) Confocal images of cardiomyocyte nuclei in control and hey2MO embryos using *Tg(myl7:nlsDsRedExpress)* at 24 hpf. (G) Bar graph showing total cardiomyocyte number at 24 hpf between control and hey2MO embryos (N=3, n=5 per condition). (H, I) Confocal images showing cardiomyocyte nuclei at 48 hpf control (H) and *hey2*^*hsc25*^ mutant embryos (I). (J) Bar graph showing cardiomyocyte number between atrium and ventricle in control and *hey2*^*hsc25*^ mutant embryos at 48hpf (N=3, n=6 per condition). Scale bars 50 μm. Error bar mean±s.e.m; ** p<0.01; *** p<0.001. A, atrium; V, ventricle; OFT, outflow tract.

### Increased proliferation of cardiac progenitors contributes to increased heart size in the absence of *hey2* function

Given the increased CM number observed in *hey2* deficient embryos from as early as 24 hpf, we explored the hypothesis that this may be due to accelerated addition of cardiac progenitors to the heart. Using whole-mount *in situ* hybridization and quantitative RT-PCR, we observed a significant increase in the expression of *nkx2.5*, *mef2cb*, *ltbp3* and *hey2* in *hey2* morphants compared to WT controls within the 24 hpf linear heart tube (Fig. 5A-J). At 48 hpf, we observed through IF staining a higher number of Mef2-positive cells at the arterial pole of the two-chambered heart (supplemental Fig. 4C-F). As we noted that the effect of *hey2* loss-of-function on CM number is evident as early as 24 hpf, we next analyzed embryos at 16.5 hpf for gene expression changes within the ALPM. Interestingly, we observed increased expression of *nkx2.5* (Fig. 5K, N and R) as well as broader expansion of *mef2cb* (Fig. 5L, O and Q) and *hey2* (Fig. 5M, P and S) expression domains. This expansion in early cardiac mesoderm gene expression was evident from as early as 12 hpf (supplemental Fig. 4A and B), implying that in the absence of *hey2* there is an expansion in the cardiac progenitor population. These results suggest that *hey2* plays an early, previously unappreciated role in restricting the size of the early cardiac progenitor population, in particular that which expresses markers of the SHF (*mef2cb, tbx1 and ltbp3*).

**Figure 5.**
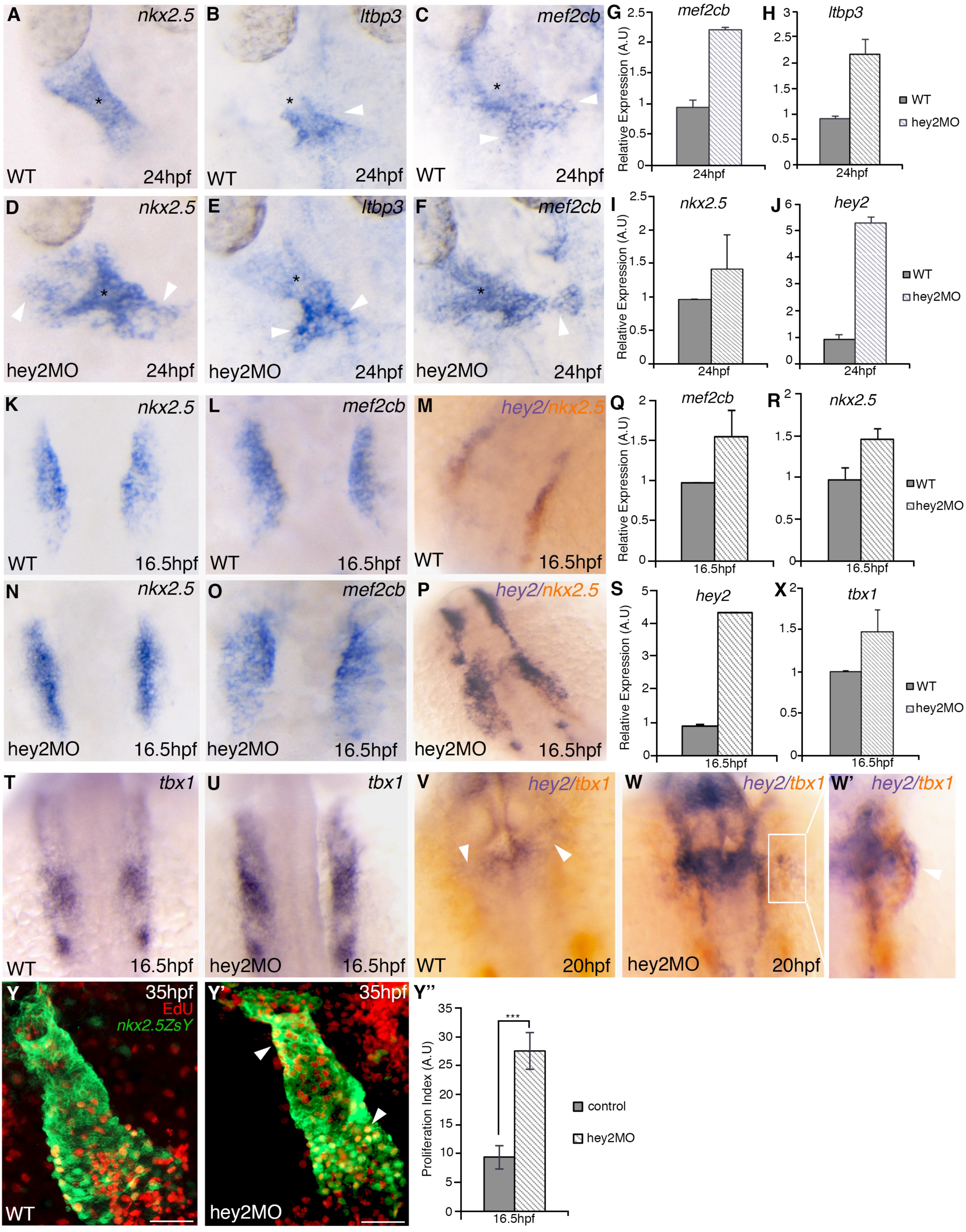
Hey negatively regulates the expression of SHF-associated genes. (A-F) Riboprobe staining for SHF markers *nkx2.5*, *ltbp3* and *mef2cb* at 24 hpf in control (A-C) and hey2MO embryos (D-F). (G-J) Quantitative RT-PCR analysis at 24 hpf comparing *mef2cb*, *ltbp3*, *nkx2.5* and *hey2* gene expression in controls to hey2MO embryos. (K-P) RNA *in situ* hybridization analysis at 16.5 hpf in control and hey2MO embryos for *nkx2.5*, *mef2cb* and combined *hey2* (blue) and *nkx2.5* (orange). (Q-S) Quantitative RT-PCR analysis for *mef2cb*, *nkx2.5* and *hey2* gene expression at 16.5 hpf. (T-W’) RNA *in situ* hybridization analysis of *tbx1* at 16.5 hpf (T and U) and co- expression of *hey2* (blue) and *tbx1* (orange) at 20 hpf (V and W, W’) in controls (T, V) and hey2MO (U, W) embryos. (W’) Enlarged view of *tbx1* and *hey2* expression within the pharyngeal mesoderm. (X) Quantitative RT-PCR analysis at 16.5 hpf for *tbx1* expression. (Y-Y’) EdU incorporation (red) in control (Y) and hey2MO (Y’) embryos expressing *Tg(nkx2.5ZsYellow)* (green). EdU positive cardiomyocytes are shown as yellow cells. (Y’’) Proliferation index between control and Hey2 morphant embryos following EdU pulse at 16.5 hpf (N=2, n=7). Error bars, mean±s.e.m; *** p<0.001.

The upregulation of SHF-associated genes observed following *hey2* loss-of-function led us to examine if there was increased proliferation of cardiac progenitors that were later added to the developing heart. Although CMs have the potential to proliferate, minimal proliferative activity has been observed within the myocardium between 24 and 48 hpf (de Pater et al., 2009). We therefore first assessed proliferation of cardiac progenitors prior to 24 hpf. These progenitors are believed to reside within pharyngeal mesoderm, in an area demarcated by *tbx1* expression, which has been shown to regulate SHF development (Buckingham et al., 2005; Chapman et al., 1996; Hami et al., 2011; Nevis et al., 2013). Tbx1 has been shown to promote proliferation of SHF progenitors and maintain the cardiac progenitor pool as myocardial accretion occurs (Nevis et al., 2013). As our initial analysis suggested that *hey2* (*epiCon21:EGFP*^*hsc28*^) is co-expressed with *Tg(nkx2.5ZsYellow)* expressing cells within the pharyngeal mesoderm (Fig. 1K’’, asterisk), we examined *tbx1* expression in *hey2* morphant embryos and found an increase in *tbx1* transcript levels at 16.5 hpf (Fig. 5T, U and X). Analysis at 20hpf revealed a partial overlap of *hey2* and *tbx1* expression domains within the pharyngeal mesoderm, with no *tbx1* expression detectable within the cardiac cone, consistent with previous reports (Fig. 5V, arrowheads; (Nevis et al., 2013). Strikingly, in the absence of *hey2* function, both *hey2* and *tbx1* transcripts were significantly up regulated within the pharyngeal mesoderm (Fig. 5W, W’ compare to V). As Tbx1 regulates SHF proliferation, we next performed an EdU incorporation assay to monitor the proliferative activity of SHF progenitors following *hey2* knockdown. *Tg(nkx2.5ZsYellow)* embryos were injected with or without *hey2* MO and pulsed with EdU at 16.5 hpf to label proliferating cells. A proliferation index was subsequently determined by comparing the number of EdU+ and EdU- cardiomyocytes at 35 hpf in control and *hey2* morphant hearts. *Hey2* morphant embryos displayed a significantly higher proliferative index as compared to controls, indicating that an increase in proliferation prior to heart tube formation contributes to enhanced CM production (Fig. 5Y-Y’’). In contrast, pulsing with EdU at 24 hpf demonstrated no significant impact of *hey2* loss on proliferation in CMs between 24-35 hpf (supplemental Fig. 4G-I). Altogether, these results suggest an early role for Hey2 in establishing the appropriate number of late-differentiating progenitors that will be added to the heart prior to heart tube formation.

### Myocardial accretion is extended in *hey2* loss-of-function embryos

Previous work has demonstrated and quantified the differentiation and addition of SHF progenitors to the heart tube between 24 and 48 hpf (de Pater et al., 2009; Hami et al., 2011; Lazic and Scott, 2011). In order to better understand the consequences of having more SHF progenitors following loss of Hey2 function, we employed *myl7:nlsKikGR* embryos to photoconvert cardiomyocytes at 24 hpf and monitor subsequent novel myocardial addition to the heart. We observed a substantial increase in the number of differentiated CMs being added to the arterial pole in *hey2* morphants (95±7.0) compared to WT controls (23.4±1.6), as shown by green only cells at the arterial pole of the 48 hpf heart (Fig. 6A-D, brackets; mean±s.e.m, n=5). The normal timing of termination for myocardial addition has been reported to be between 36 and 48 hpf (Jahangiri et al., 2016; Lazic and Scott, 2011). Due to the increase in cell addition between 24 and 48 hpf, we wondered whether myocardial accretion was extended temporally in the absence of Hey2. By photoconverting embryos at 48 hpf, with subsequent imaging at 60 hpf, we noted SHF-mediated accretion in control embryos was minimal, consistent with previous findings (Fig. 6F and H, 11.9±0.7, mean±s.e.m, n=7; (de Pater et al., 2009; Jahangiri et al., 2016). In *hey2* deficient embryos a significantly increased cell addition was evident beyond 48 hpf (Fig. 6G and H, 20.3±1.3, mean±s.e.m, n=7). This highlights that in the absence of Hey2, the window of myocardial accretion from the late-differentiating progenitor population becomes extended. Following cell addition to the heart, a subpopulation of SHF progenitors has shown to gives rise to the OFT, a structure containing both myocardium, for lengthening the distal cardiac portion, as well as smooth muscle at the myocardial-arterial junction (Choi et al., 2013; Hami et al., 2011; Waldo et al., 2005; Zeng and Yelon, 2014; Zhou et al., 2011). Given that loss of *hey2* resulted in increased proliferation and extended the window of myocardial accretion by cardiac progenitors, we wondered what effect this may have on other lineages of the SHF, in particular the OFT smooth muscle. Via incubation of wild type and *hey2*^*hsc25*^ mutant embryos in DAF-2DA, a compound which specifically labels smooth muscle of the OFT (Grimes et al., 2006), we found that *hey2*^*hsc25*^ mutants had a significantly longer OFT than that of sibling controls (Fig. 6I-K). Together this demonstrates that in the absence of Hey2, late-differentiating progenitors are subjected to an extended window of accretion following an increase in proliferative activity, which ultimately leads to an expansion in SHF-derived structures.

**Figure 6.**
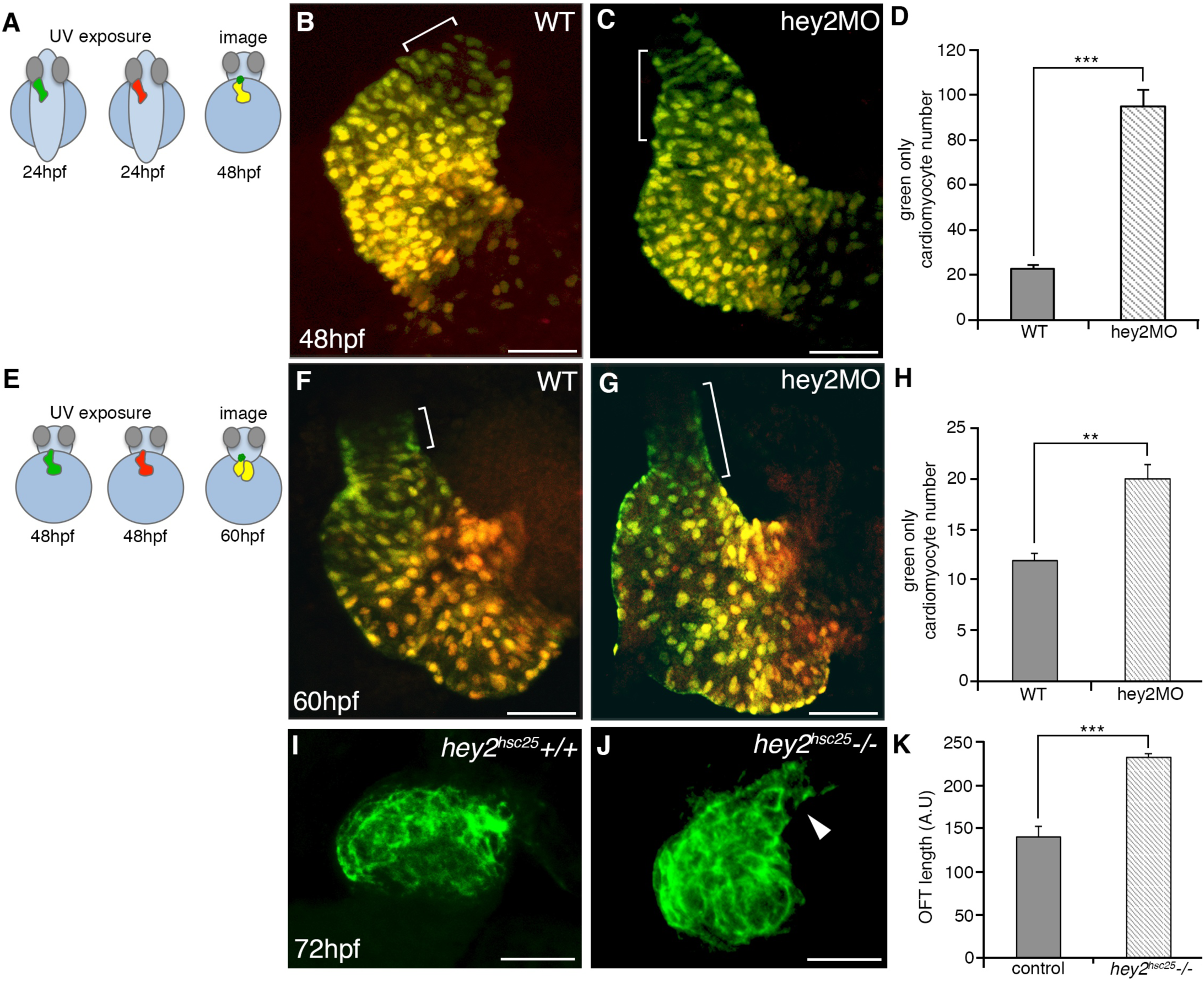
Hey restricts SHF cell addition to the developing heart. (A) Schematic representation of cardiomyocyte photoconversion assay between 24 and 48 hpf. (B-C) Confocal imaging of control and hey2MO *Tg(myl7:nlsKikGR)* embryos. Brackets highlight green only SHF-derived cardiomyocytes. (D) Bar graph showing a significant increase in green-only cardiomyocytes at 48 hpf in control compared to Hey2 morphant embryos (N=3, n=7). (E) Schematic representation of photoconversion assay between 48 and 60 hpf. (F-G). Confocal imaging of control and hey2MO *Tg(myl7:nlsKiKGR)* embryos showing an increase in SHF-derived green only cardiomyocytes in morphants (G) compared to controls (F). (H) Bar graph showing mean values of green-only cells between 48 and 60 hpf (N=3, n=7). (I-K) DAF2-DA labeling of the OFT smooth muscle in control and *hey2*^*hsc25*^ mutant embryos at 72 hpf. (K) Bar graph represents mean OFT lengths in control and *hey2*^*hsc25*^ mutant (N=2, n=11 per condition). Error bars mean±s.e.m; ^**^ p<0.01; *** p<0.001. Scale bars 50μm.

### *hey2* acts cell autonomously to regulate SHF addition to the heart at the expense of the FHF population

We next employed a transplantation approach to examine the cell autonomy of Hey2 activity in cardiac progenitors. To accomplish this, *Tg(myl7:nlsKikGR)* donor embryos, either WT or injected with *hey2* MO, were used. Donor cells at 4 hpf (50% epiboly) were transplanted to the margin of WT host embryos (Fig. 7A), an approach that has been shown to result in cardiac contribution of donor cells (Scott et al., 2007; Stainier et al., 1993). Transplant embryos were photoconverted at either 24 or 48 hpf and imaged at either 48 or 60 hpf (Fig. 7B and F, respectively). From our results we observed that *Hey2* morphant donor *Tg*(*myl7:nlsKikGR)* cells displayed a significant increase in SHF contribution (shown by the ratio of green:yellow cardiomyocytes per embryo), as compared to early myocardial addition, between 24 to 48 hpf (Fig. 7C-E). The same result was observed between 48 to 60 hpf, with Hey2 deficient *myl7:nlsKikGR* donors contributing significantly more green only cardiomyocytes than controls (Fig. 7F-I). While the ratio in numbers of late versus early differentiated cardiomyocytes per heart from *hey2* donor cells was consistently increased at both 24-48 and 48-60 hpf, an analysis of cell numbers for each category revealed a bias in progenitor populations. From 24-48 hpf, the increased late versus early CM addition ratio observed in *hey2* morphants was due to a decreased propensity for donor cells to contribute early to the heart, evident by a significant decrease in total cell number of yellow CMs in controls compared to morphant embryos (Fig. 7J). However, when comparing total number of late differentiating CMs, no statistical significance was observed (Fig. 7J). In contrast, the higher late versus early addition ratio observed at 48 hpf was due to a significantly higher amount of late (post 48 hpf) CM addition, with a relatively equivalent amount of early (pre 48 hpf) CM addition as noted by no significant change in early CM number between control and *hey2* morphant embryos (Fig. 7K). These results demonstrate a cell-autonomous function for Hey2, in presumptive cardiac progenitors, that delays their addition to the heart as cardiomyocytes.

**Figure 7.**
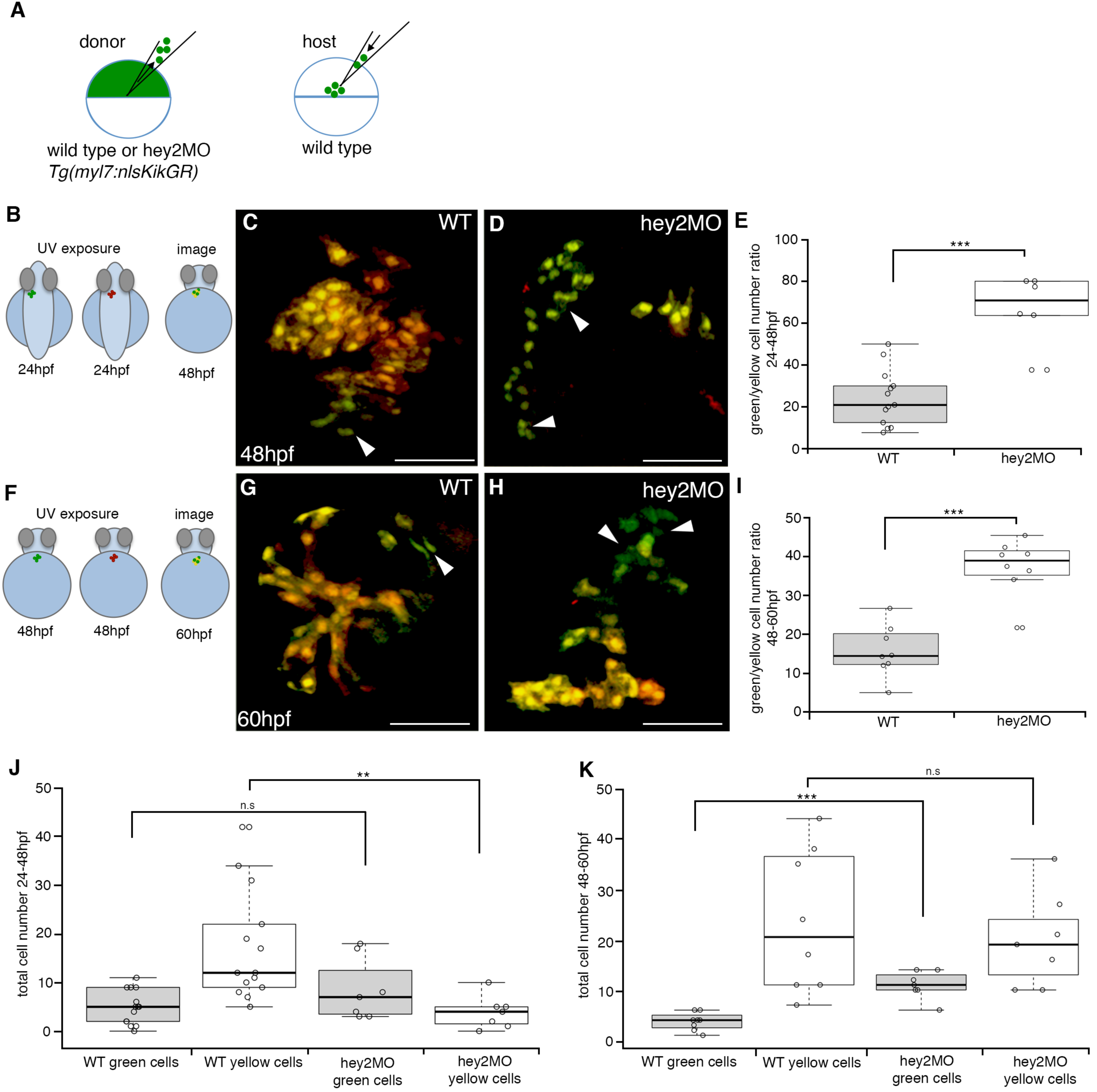
Hey2 functions cell-autonomously to inhibit SHF progenitor contribution to the developing heart. (A) Schematic representation of transplantation strategy. (B-D) photoconversion *Tg(myl7:nlsKiKGR)* at 24 hpf, imaged at 48 hpf in control (C) and hey2MO (D) transplanted embryos. (E) Boxplot analysis demarking the percentage ratio at 24-48 hpf between green and yellow transplanted cardiomyocytes in control and hey2MO embryos (n=13, control and n=7, hey2MO). (F-H) photoconversion at 48 hpf, imaged at 60 hpf of *Tg(myl7:nlsKiKGR)* control (G) and hey2MO (H) transplants. (I) Boxplot analysis demarking the percentage ratio at 48-60 hpf between green and yellow transplanted cardiomyocytes in control and hey2MO embryos (n=8, control and n=9, hey2MO). (J-K) Boxplot analysis displaying total cell number of transplanted *Tg(myl7:nlsKikGR)* control and hey2MO cardiomyocytes at 24-48 hpf (J; n=12 WT green, n=15 WT yellow; n=7 hey2MO green, n=7 hey2MO yellow) and 48-60 hpf (K; n=8 WT green, n=8 WT yellow; n=7 hey2MO green, n=7 hey2MO yellow). Green cells refer to SHF contribution; yellow cells refer to FHF contribution. Error bars, mean±s.e.m; ^**^ p<0.01, *** p<0.001; n.s, no significant difference. Scale bars 50μm.

## DISCUSSION

Our work demonstrates a novel role for the bHLH factor Hey2 in regulating the size of the cardiac progenitor pool and the timing of the contribution of late-differentiating cardiac progenitors to the zebrafish heart. As shown by fate mapping (Camp et al., 2012; Mjaatvedt et al., 2001) and lineage tracing (Cai et al., 2003; Meilhac et al., 2003) approaches, the vertebrate heart is made via the temporally distinct addition of at least two populations of cardiac progenitors. A recent study using live imaging of cell lineage tracing and differentiation status suggests that in mouse a discrete temporal lag can be observed between the first and second waves of differentiation that form the heart (Ivanovitch et al., 2017). Whether the cardiac progenitor pools that are added early and late to the heart, termed the FHF and SHF, represent molecularly distinct populations remains an open question. While FHF- and SHF-restricted cells can be identified at early gastrulation stages in mouse by clonal analysis (Devine et al., 2014; Lescroart et al., 2010), this may reflect the result of distinct migratory paths and signaling milieus experienced by cells during gastrulation.

How the relative size of cardiac progenitor pool(s), the timing of their differentiation and the extent to which they are added to various components of the heart all remain largely unknown. Our work has uncovered a cell autonomous function for Hey2 to restrain proliferation of cardiac progenitors prior to their addition to the heart. Our expression analyses suggest that *hey2* co-localizes with previously identified SHF-associated genes including *isl2b* and *mef2cb* (Figure 1; (Lazic and Scott, 2011; Witzel et al., 2017) and is in agreement with fate mapping work that has shown SHF progenitors to reside in an anteromedial position within the ALPM (Hami et al., 2011). Importantly, we and others have shown that *hey2* expression is regulated not by a canonical Notch signaling pathway, but by RA/FGF signaling (Sorrell and Waxman, 2011). This is consistent with the Notch-independent, FGF-mediated expression of *Hey2* in other developmental contexts (Doetzlhofer et al., 2009). Previous reports have highlighted that the cardiac malformations found in animals lacking *Hey2* function resemble common human congenital heart defects including ventricular septal defects, tetralogy of fallot and tricuspid atresia (Donovan et al., 2002). Coupled with the expression of *hey2* during cardiac development, the effects of *hey2* loss on late myocardial addition to the zebrafish heart suggest a mechanism where Hey2 acts specifically in SHF progenitors. While we did not observe appreciable effects on FHF-associated markers in our study, this data is difficult to interpret, as bona fide FHF- and SHF-specific markers that distinguish these populations are poorly characterized. Our transplant assays strongly suggest that cardiac contribution is at the very least delayed with the loss of *hey2*. The observed phenotypes may reflect: 1) a shift in the balance between FHF and SHF progenitor proliferation, favouring the SHF pool; 2) alteration in the timing of cardiac progenitor differentiation; 3) changes in the number of progenitors allocated to the FHF and SHF pools; or 4) a FHF/SHF-agnostic role for Hey2 in cardiac progenitor proliferation and differentiation. This is a critical question that will require further study, but that may provide insight into the diversity of early cardiac progenitors.

It is important to note that the function of *hey2* in zebrafish cardiogenesis has been previously addressed in an elegant study (Jia et al., 2007). However, while the overall “large heart” phenotype observed is shared between our *hey2*^*hsc25*^ and the *grl*^*m145*^ mutants, our data suggests a role for *hey2* prior to 24 hpf, in cardiac progenitors, that subsequently impacts cardiac development. In contrast, Jia and colleagues reported that *grl* had minimal affect during this time, with *grl*^*m145*^ mutants having comparable cardiomyocyte numbers to controls at 24 hpf. This discrepancy may reflect the nature of the *hey2* alleles used, with the *hey2*^*hsc25*^ allele being, we believe, a true null. As the SHF and late myocardial addition in zebrafish had not been described at the time of the prior study, this would have also affected the interpretation of the results. This highlights the fact that Hey2 likely acts at multiple steps of heart development.

The myocardium of the AVC is important for the development of the AV cushion and AV node, both derivatives of the SHF (Kelly, 2012). In zebrafish, *bmp4* and *tbx2b* are expressed in the AV myocardium, and play critical roles in the establishment of AVC identity (Ma et al., 2005; Zhang and Bradley, 1996). Here we demonstrate the importance of Hey2 in AVC development. Although previous work revealed that the *grl*^*m145*^ mutant shows an ectopic expansion of *bmp4* expression at 48hpf, little change was observed in *tbx2b* transcripts (Rutenberg et al., 2006). However our mutant *hey2*^*hsc25*^ allele revealed upregulation in both *bmp4* and *tbx2b*, thus providing insight into a potential regulatory network by which *hey2* expression in ventricular myocardium constrains *bmp4* expression to the AVC, which in turn activates *tbx2b* expression (supplemental Fig. 4L). In the absence of functional Hey2, this repression is absent, resulting AVC-specific genes expanding their expression domains into the cardiac chambers (supplemental Fig. 4M).

Our data suggest multiple roles for Hey2 in cardiogenesis, first acting to restrain cardiac progenitor proliferation (and potentially affecting diversification of FHF and SHF progenitors), and later affecting the timing of when SHF progenitors are added to the developing heart. Hey2 may therefore be an intrinsic regulator of the extent and differentiation of the cardiac progenitor pool, possibly acting as a readout of extrinsic (RA and FGF) niche signals. It remains to be determined if this strictly reflects a role for Hey2 in SHF progenitors, or if Hey2 plays a broader role in all cardiac progenitors. To address these questions, lineage tracing approaches will be required, for which the novel *hey2* enhancer transgenics we have uncovered will be of great utility. Given the cell autonomous function of Hey2, identifying its transcriptional targets will also be of great interest. Given the role of Hey2 in restraining cardiac progenitor proliferation, its down-regulation by RA, and the known role of epicardial RA signaling in zebrafish heart regeneration (Kikuchi et al., 2011), a potential role for Hey2 in regeneration should also be investigated. Dissecting how Hey2 regulates cardiac development will help address key unanswered questions with respect to the regulatory mechanisms that coordinate the size and differentiation timing of cardiac progenitors to allow for proper heart development to proceed.

## ACKNOWLEDGMENTS

We would like to thank Angela Morley and Allen Ng for expert fish care and maintenance. Neil Chi kindly provided the *myl7:EGFP* and Caroline and Geoffrey Burns the *nkx2.5ZsYellow* transgenic lines. NG and XY were kindly supported by a Labatt Family Heart Centre Philip Witchel post-doctoral fellowship and a Hospital for Sick Children Restracomp studentship, respectively. Research funding was generously provided by the Heart and Stroke Foundation of Canada (to ICS and MDW, Grant-in-Aid G-16-00013798), the Natural Sciences and Engineering Research Council of Canada (to ICS, RGPIN 341545-12) and the Canadian Institutes of Health Research (Operating Grant MOP-123223 (to ICS) and Project Grant PJT - 153343 (to ICS and MDW)).

## FIGURE LEGENDS

**Supplemental Figure 1:**
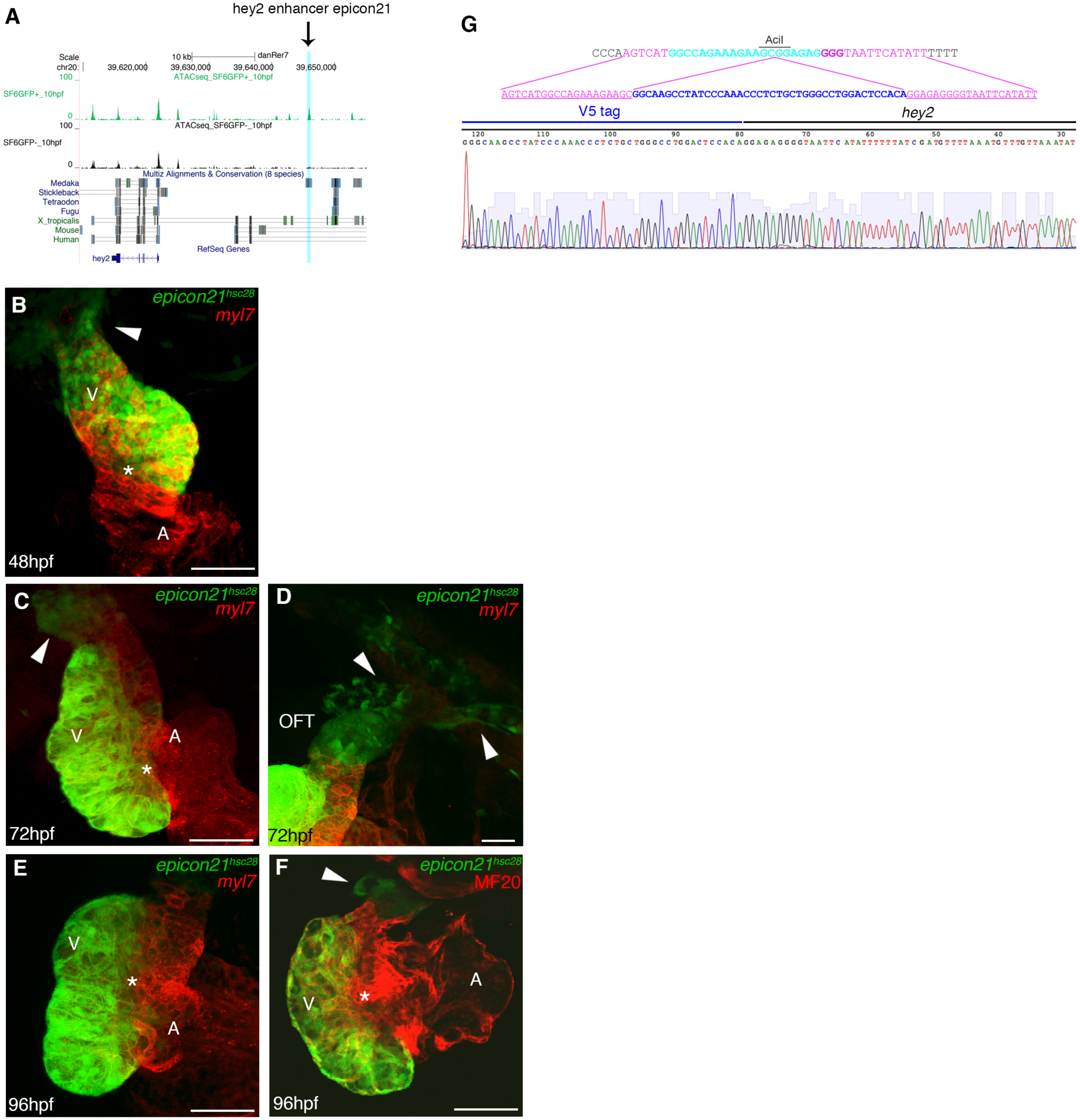
related to Figure 1. (A) Schematic representation of hey2 enhancer (epigenetic conserved region 21) located 24kb upstream of *hey2* locus used to create *Tg(epiCon21:EGFP)*^*hsc28*^. (B-F) Confocal microscopy of *Tg(epiCon21:EGFP)*^*hsc28*^ at 48 hpf (B), 72 hpf (C, D) and 96 hpf (E, F) against *Tg(myl7:nlsDsRedExpress).* (G) Schematic methodology of internal epitope tagging of V5 into *hey2* locus with corresponding sequence of tag-specific PCR fragments. Scale bars 50μm. OFT, outflow tract; V, ventricle and A, atrium; Asterisk, atrioventricular canal. Arrowheads mark the *epiCon21+* OFT region.

**Supplemental Figure 2:**
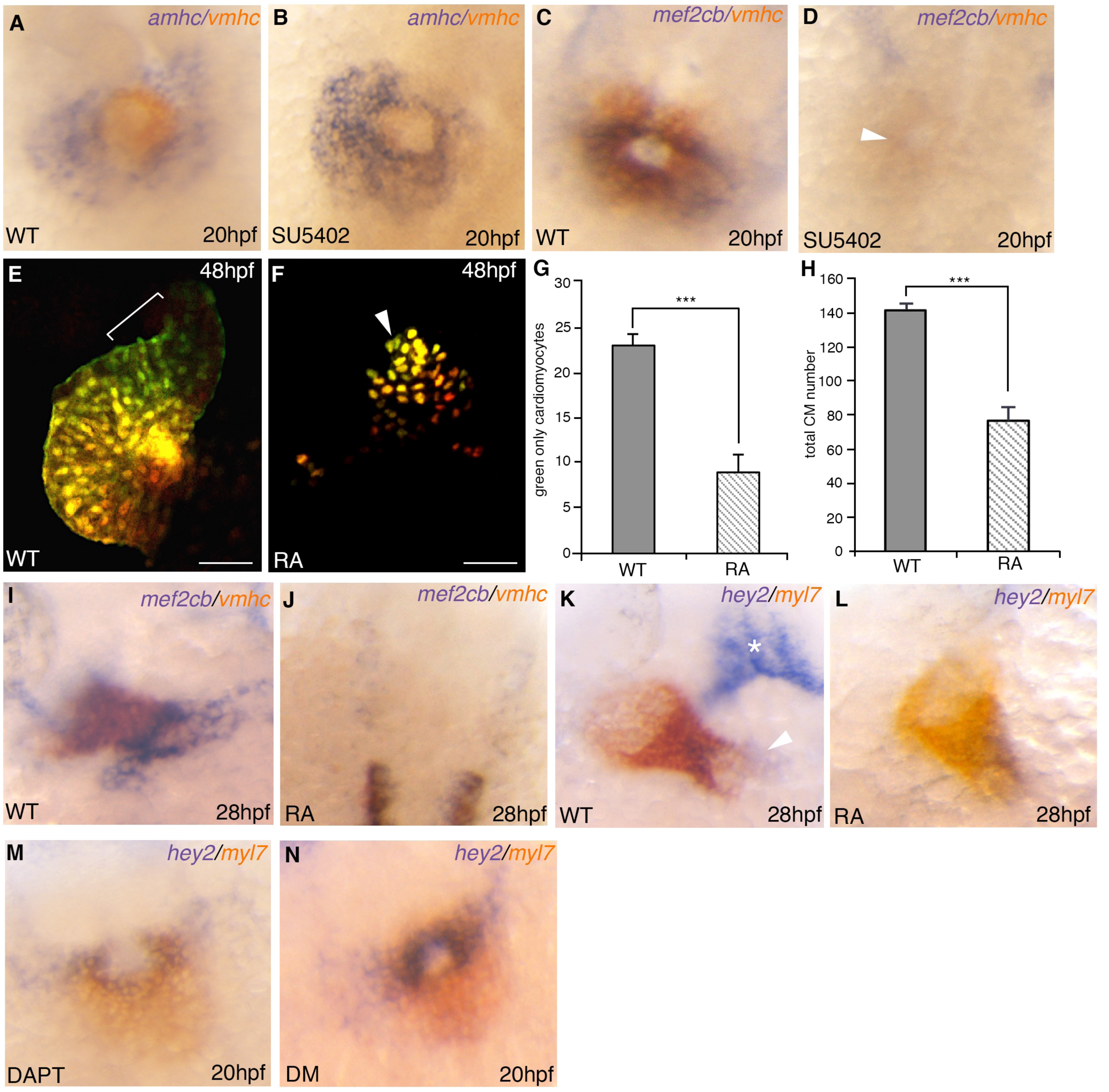
related to Figure 2. (A-D) double *in situ* hybridization analysis for *amhc* (blue), *vmhc* (orange) and *mef2cb* (blue) at 20 hpf following treatment with 10μM SU5402 from 16.5 hpf. (E-F) Confocal imaging at 48 hpf of photoconverted *Tg(myl7:nlsKikGR)* embryos at 24 hpf following treatment with 0.1uM RA at 50% epiboly. (G-H) Bar graph analysis showing the number of green-only SHF cell addition by 48 hpf (G) and total ventricular cell number (H). N=2, n=5 per condition. Error bars mean±sem; *** p<0.001. Scale bars 50μm.

**Supplemental Figure 3:**
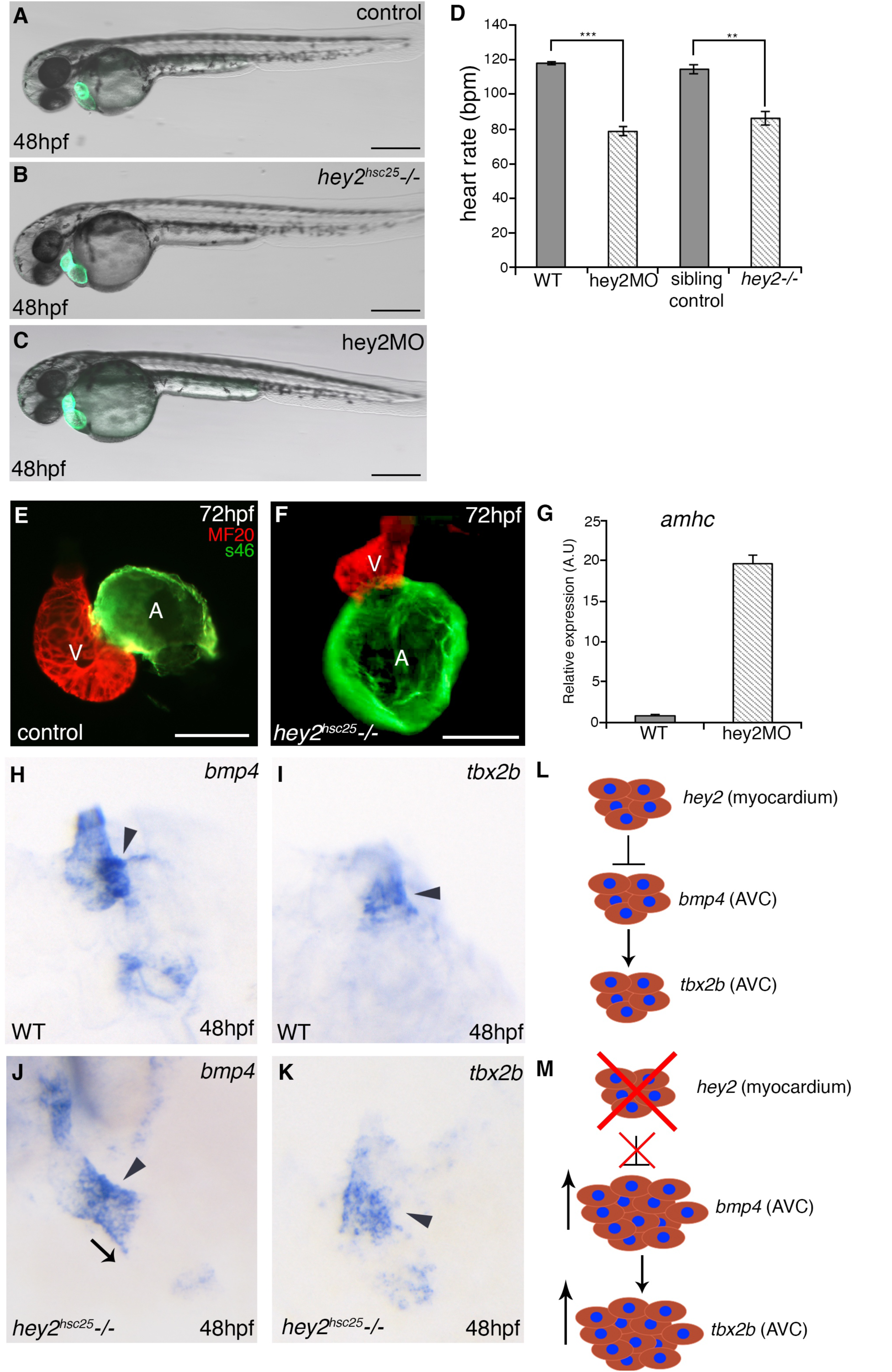
related to Figure 3. (A-C) Bright-field images of *Tg(myl7:EGFP*) in control (A), *hey2*^*hsc25*^ mutant (B) and hey2MO (C) embryos at 48 hpf. (D) Heart rate analysis represented as beats per minute (bpm) at 48hpf in control, hey2MO and *hey2*^*hsc25*^ mutant embryos (N=3, n=4). (E-F) MF20/S46 immunofluorescence imaging at 72 hpf in control (E) and *hey2*^*hsc25*^ mutant embryos (F). (G) Quantitative RT-PCR analysis for *amhc* gene expression in control and hey2MO embryos at 48 hpf (gene expression normalized to β-actin, fold difference relative to control; N=3, n=3). (H-K) Riboprobe staining for *bmp4* (H andn J) and *tbx2b* (I and K) at 48 hpf in control and *hey2*^*hsc25*^ mutant embryos. (L-M) Schematic representation showing boundary constraints of *bmp4* and *tbx2b* genes expression within the ventricular myocardium and AVC in the presence (L) and absence (M) of *hey2*. *** p<0.001, ^**^ p<0.01, error bars mean±s.e.m. Scale bars 50μm.

**Supplemental Figure 4:**
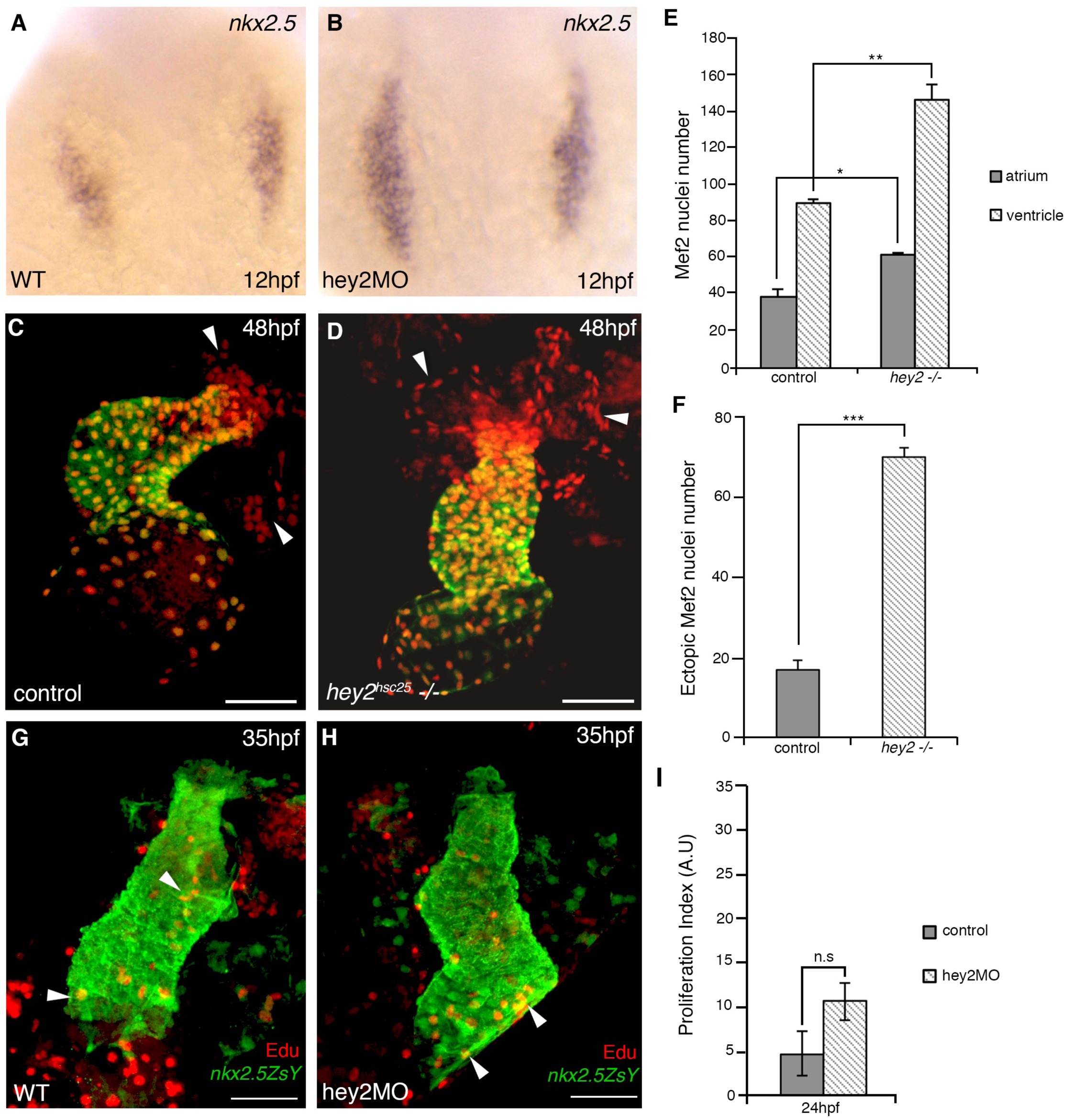
related to Figure 5. (A-B) Riboprobe staining for *nkx2.5* transcripts at 12 hpf in WT and hey2MO embryos. Expanded expression of *nkx2.5* was observed in morphants (B) compared to controls (A). (C-D) Confocal imaging of Mef2 in *Tg(myl7:EGFP)* control embryos (C) and *hey2*^*hsc25*^ mutants (D). (E-F) Bar graph analysis displaying total number of Mef2+ cells in the heart proper (E) and adjacent to the arterial pole (F, arrowheads) between controls and mutants at 48 hpf (N=3, n=4). (G-H) Confocal imaging of EdU incorporation assay. Embryos were pulsed at 24 hpf and imaged at 35 hpf. (I) Bar graph showing the proliferative index of Edu+/*nkx2.5+* cardiomyocytes between controls and hey2 morphants when pulsed at 24 hpf (N=2, n=4). No significant difference in proliferation was observed. Error bars mean±sem; ^*^ p<0.05, ^**^ p<0.01, *** p<0.001; n.s, no significance. Scale bars 50μm.

